# Nicotinic modulation insecticides act on diverse receptor subtypes with distinct subunit compositions

**DOI:** 10.1101/2021.11.03.467052

**Authors:** Wanjun Lu, Zhihan Liu, Xinyu Fan, Xinzhong Zhang, Xiaomu Qiao, Jia Huang

## Abstract

Insect nicotinic acetylcholine receptors (nAChRs) are ligand gated ion channels mainly expressed in the central nervous system of insects. They are the directed targets of nicotinic modulation insecticides including neonicotinoids, the most widely used insecticides in the world. However, the resistance development from pests and the negative impacts on the pollinators affect their applications and create demand for the alternatives. Thus, it is very important to understand the mode of action of these insecticides at the molecular level, which is actually unclear for more than 30 years. In this study, we systematically examined the susceptibility of ten *Drosophila melanogaster* nAChR subunits mutants against eleven nicotinic modulation insecticides. Our results showed that there are several subtypes of nAChRs with distinct subunits compositions that are responsible for the toxicity of different insecticides, respectively. At least three of them are the major molecular targets of seven structurally similar neonicotinoids *in vivo*. On the other hand, the spinosyns may exclusively act on the α6 homomeric nAChR but not any other heteromeric pentamers. Behavioral assays using thermogenetic tools further confirmed the bioassay results and support the idea that receptor activation rather than inhibition leads to the insecticidal effects of neonicotinoids. The present findings reveal native nAChR subunit interactions with various insecticides and have important implications for resistance management and the development of novel insecticides targeting this important ion channels.

**Author Summary:** The neonicotinoids and spinosyns make up about 27% of the insecticides by world market value. Novel insecticides like sulfoxaflor, flupyradifurone and triflumezopyrim are developed as alternatives due to the negative effects of neonicotinoids on pollinators. Although all act via insect nicotinic acetylcholine receptors, the mode of action is unclear. Our work shows that these insecticides act on diverse receptor subtypes with distinct subunit compositions. This finding could lead to the development of more selective insecticides to control pests with minimal effects on beneficial insects.

## Introduction

Chemical insecticides have been wildly used to control pests in agriculture, horticulture, forestry, homes and cities. They have also played a vital role in preventing the spread of human and animal vector-borne diseases. However, Insecticide resistance is a serious worldwide problem for invertebrate pest control, with more than 600 different insect and mite species having become resistant to at least one insecticide. In addition, there is at least one documented case of resistance for more than 335 insecticides/acaricides[1]. Therefore, there is great demand for effective insecticide resistance management (IRM) and development of new pest control compounds. To address both issues, we need to know the mode of action of insecticides: the process of how an insecticide works at a molecular level [2].

A complete understanding of the mode of action of an insecticide requires knowledge of how it affects a specific target site within an organism. Although most insecticides have multiple biological effects, toxicity is usually attributed to a single major effect. For many insecticides, however, the exact molecular targets remain elusive. In order to ascribe whether a candidate protein is indeed the target for an insecticidal effect *in vivo*, it is not sufficient to demonstrate an *in vitro* biochemical interaction between an insecticide and a protein. Genetic evidence demonstrating an effect due to mutation of the candidate target is critical before it is possible to conclude that a given protein is the target of an insecticide.

The neonicotinoids (acetamiprid, clothianidin, dinotefuran, imidacloprid, nitenpyram, thiacloprid, and thiamethoxam) are remarkably effective to control agricultural pests, ectoparasites and arthropod vectors [3]. They are taken up by the roots or leaves and translocated to all parts of the plant due to high systemic activity, making them effectively toxic to wide range of sap-feeding and foliar feeding insects. Thus, neonicotinoids account for 24% of the global insecticide market, the largest market share of all chemical classes[1]. They act selectively on insect nicotinic acetylcholine receptors (nAChR) as agonists compared with the mammalian-selective nicotine. The spinosyns are a naturally derived, unique family of macrocyclic lactones which act on insect nAChR in an allosteric fashion. Besides, the sulfoximine sulfoxaflor, butenolide flupyradifurone and mesoionic triflumezopyrim are three newly developed insecticides which are also nAChR competitive modulators [4]. It is expected that the market of all above nAChR targeting insecticides which show excellent insect to mammalian selectivity, will continue to grow. However, the molecular targets of neonicotinoids and other nAChR modulators remain unclear, mainly because we do not know the structure and assembly of native nAChRs in insects [5].

The cation-selective nAChRs are members of the Cys-loop ligand gated ion channel superfamily responsible for rapid excitatory neurotransmission. The functional nAChRs are homo- or heteromeric pentamers of structurally related subunits arranged around a central ion-conducting pore[6]. Each subunits has a extracellular N-terminal domain which contains six distinct regions (loops A–F) involved in ligand binding, four C-terminal transmembrane segments (TM1–TM4) and an intracellular loop between TM3 and TM4. nAChRs are divided into α-subunits possessing two adjacent cystine residues in loop C, while those subunits without this motif are termed non-α subunits. In vertebrates, 17 nAChR subunits have been identified, which can co-assemble to generate a diverse family of nAChR subtypes with different pharmacological properties and physiological functions. Insects have fewer nAChR subunits (10–12 subunits) according to the available genome data. Although co-immunoprecipitation studies have indicated potential associations of several subunits, the exact subunits composition of native insect nAChRs remains unknown[5]. Unlike the vertebrate counterparts, heterologous expression of genuine arthropod α and β subunits has not been successful until recently two groups found that three ancillary proteins are essential for robust expression of arthropod nAChR heteromers [7, 8]. Thus for a long time, researchers used hybrid receptors with insect α subunits and mammalian/avian β subunits to study the interaction of insecticides and receptors. Such alternatives may not faithfully reflect all features of the native nAChRs [9].

In this study, we systematically examined the effects of total ten (seven α and three β) *Drosophila melanogaster* subunit mutants against eleven different nAChR targeting insecticides. We found that there are multiply subtypes of receptors with distinct subunits compositions which are responsible for the toxicity of different insecticides, respectively. Artificial activation/inhibition of subunit-expressing neurons also mimicked insecticides poisoning symptoms in pests. The elucidation of molecular targets of these economically important agrochemicals and the assembly of native nAChRs will be very helpful for resistance management and ecotoxicological evaluation on beneficial insects like predators and pollinators.

## Results

### Generation of nAChRβ1^R81T^ mutant

We got all 10 nAChRs knock-out mutants from Yi Rao’s lab and found that KO of α4 and β1 was homozygous lethal. Thus we used a point mutation (T227M) allele of α4 (*redeye*, *rye*) in bioassays, which is a dominant-negative mutation to cause reduced sleep phenotype in flies [10]. An R81T mutation of the nAChR β1 was found in neonicotinoids-resistant peach aphids and later in cotton aphids [11, 12], so we introduced the homologous mutation into the β1 of *Drosophila melanogaster* with CRISPR-Cas9–mediated homology-directed repair (HDR). The design of gRNA target site and HDR template was shown and the screen of successful R81T knock-in was accreted under imidacloprid selection pressure and confirmed by direct DNA sequencing (Figure 1 and S1).

**Figure 1.**
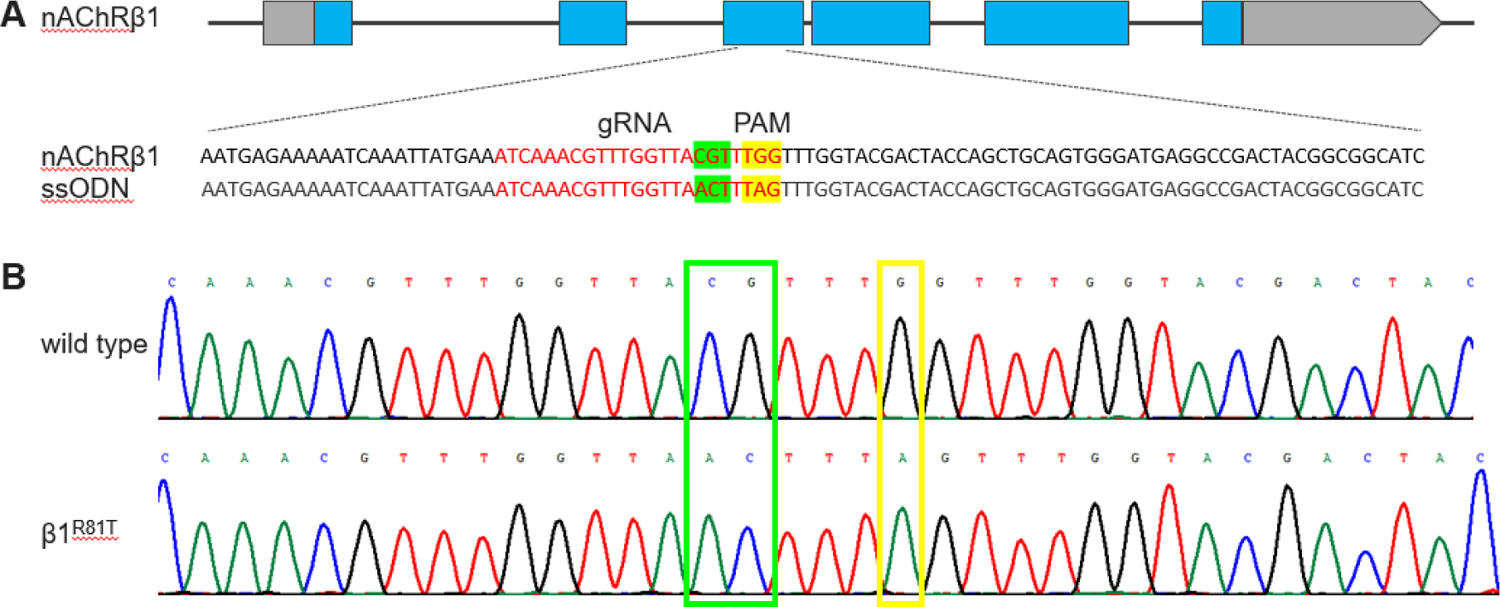
Generation of the nAChRβ1^R81T^ mutant by CRISPR/Cas9 genome editing. (A) Schematic of the nAChRβ1 locus and the sequence of donor construct. The boxes represent exons and the coding regions are shown in blue. The gRNA sequence is indicated in red and the coden for amino acid substition (CGT to ACT) is highlighted in green. One synonymous mutation (G to A) is also introduced in the PAM region (in yellow) to prevent the re-cleavage from Cas9 after successful integration. (B) The sequence comparison between wild type and point mutation flies. The nucleotides replaced are highlighted in green and yellow boxex.

### nAChR mutants showed distinct resistance to multiple insecticides

We testeded the effects of 10 nAChR mutants and some heterozygous mutants against 11 insecticides (Figure 2 and Table S1-11). The α1 mutant showed moderate levels of resistance to imidacloprid, thiacloprid, acetamiprid and triflumezopyrim, the LC_50_ resistance ratio (RR) is about 13.5 - 88.0. Its heterozygous mutant also showed low levels of resistance to these insecticides. Besides, it showed a low but statistically significant increases of RR (2.7 - 3.7) to thiamethoxam, clothianidine, dinotefuran and nitenpyram. The α2 mutant also showed similar levels of resistance (17.2 - 48.5 in the terms of RR) to imidacloprid, thiacloprid and triflumezopyrim. For the α3 mutant, it showed small RR increases (2.7 - 5.5) to thiamethoxam, clothianidine, dinotefuran, nitenpyram, sulfoxaflor and flupyradifurone. The α4, α5, α6, α7 and β3 mutants are sensitive to almost all the tested insecticides, the obvious exception is that the α6 homozygous mutant is resistant to spinetoram with a RR of 42.8 but the heterozygous mutant is close to the wild type (RR 1.2). The β1 mutant exhibited medium to high resistance to all insecticides (23.9 - 398.3 in the terms of RR) except spinetoram, and its heterozygous mutant showed small increases of RR for most insecticides. The resistance profile of β2 mutant is similar to that of α1 mutant, with 13.0 - 84.3 folds RR increases to imidacloprid, thiacloprid, acetamiprid and triflumezopyrim.

**Figure 2.**
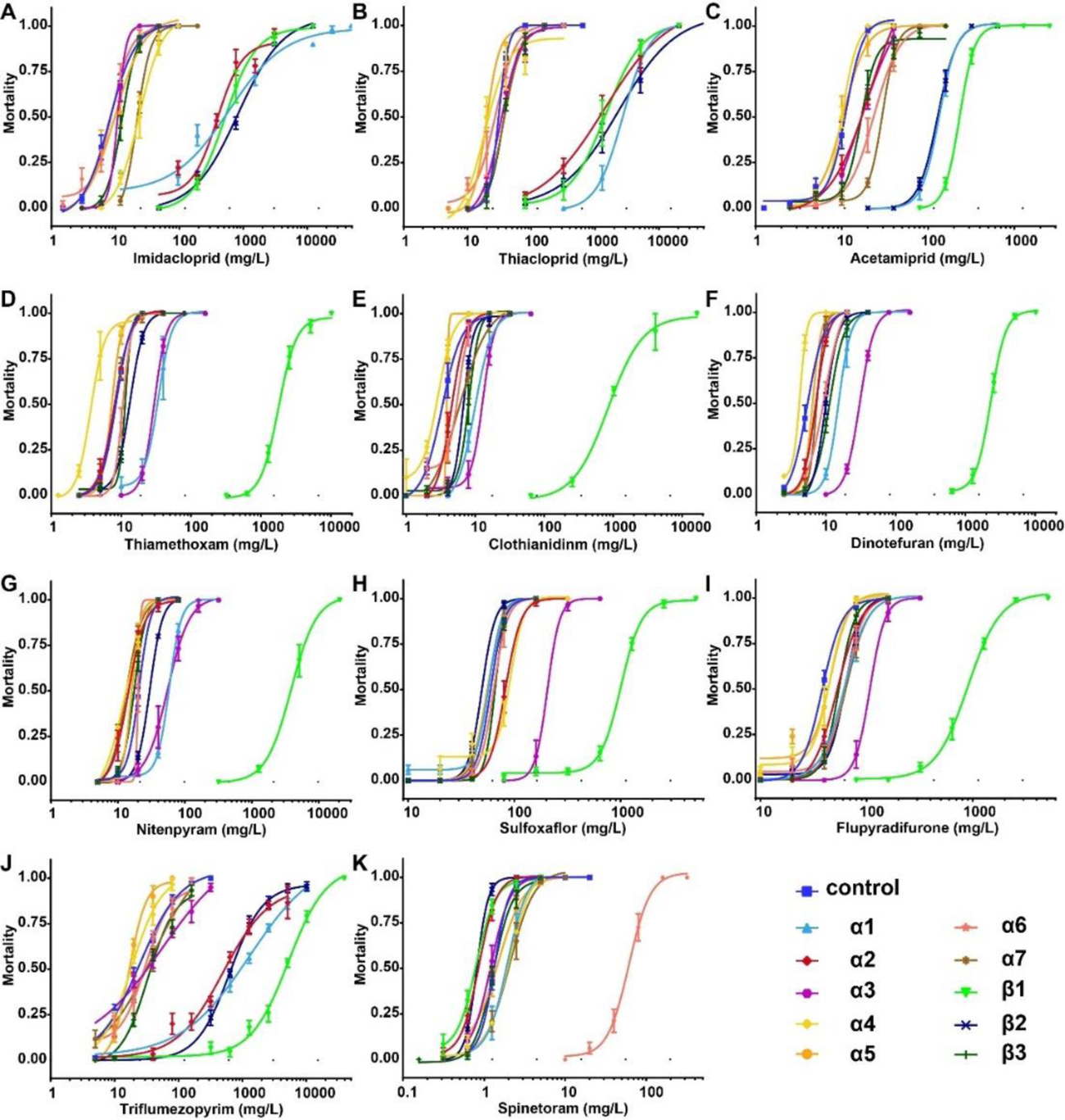
Non-linear Log-dose mortality data for tested insecticides against ten *Drosophila* nAChR homozygous mutants including eight null alleles and two point mutation alleles (α4^T227M^ and β1^R81T^). Mortality (0-1 means 0-100% in terms of percentage) of control and mutant female adults after 48 hour exposure to increasing concentrations of insecticides. Error bars represent standard deviations.

Both α1 and β1 mutants showed variable resistance to multiply insecticides, thus we generated a α1/β1 double mutant with recombination. However, the eggs laid by this combined mutant can not hatch for further experiments. A recent paper also generated a β1 R81T *Drsosophila* and found that it has serious defects in reproduction and locomotion [13], however, the β1 mutant we made here did not show any significant fitness cost (Figure S2). The sequences of α5, α6 and α7 are very close and show high similarity to the vertebrate nAChR α7 subunit. They can also form heteromecic ion channels in vitro with different combinations like α5/α6, α5/α7 and α5/α6/α7 [14]. Since only the α6 mutant showed resistance to spinetoram, we then wonder whether there is a genetic redundancy among these evolutionarily conserved gene. However, the α5/α7 double mutant was still sensitive to spinetoram (Table S11), indicating that the α6 homomeric channel could be the solo target for spinosyns.

### Hyperactivating/silencing *nAChR*-expressing neurons mimics insecticides poisoning symptoms

The way insects react when they are exposed to neonicotinoids, sulfoxaflor, flupyradifurone and spinosyns are similar. The early-onset behaviors including hyperactivity, convulsion, uncoordinated movements, leg extension and tremors. At higher doses, these excitatory symptoms can induce severe tremors and complete paralysis that lead to death [15–17]. We then wondered whether artificial activation of *nAChR-expressing* neurons wound induce insecticides-like poisoning symptoms. Thus, we used the thermosensitive cation channel *Drosophila* TRPA1 to acutely hyper-stimulate these neurons with all available *nAChR* KI-Gal4 strains [18]. We found that expressing *trpA1* in *nAChRα1^2A-GAL4^*-*nAChRα2^2A-GAL4^*, *nAChRα3^2A-GAL4^*, *nAChRα6^2A-GAL4^ and nAChRβ2^2A-GAL4^* neurons strongly induced hyperactivity behavior at 32 ℃, and eventually led to paralysis (Figure 3A, Video 1), which is similar to the above-mentioned symptoms. However, activation of *nAChRβ3^2A-GAL4^* neurons did not show any behavioral defects. These results parallel the above bioassay data that the deletion of α1, α2, α3, α6 and β2 caused medium to high resistances to these insecticides respectively. Therefore, thermogenetic activation of some *nAChR-* expressing neurons in a short time window phenocopies the action of insecticides in target pests, which demonstrates that *in vivo* pharmacological activation of these subunits-containing nAChRs leads to toxicity and finally death.

**Figure 3.**
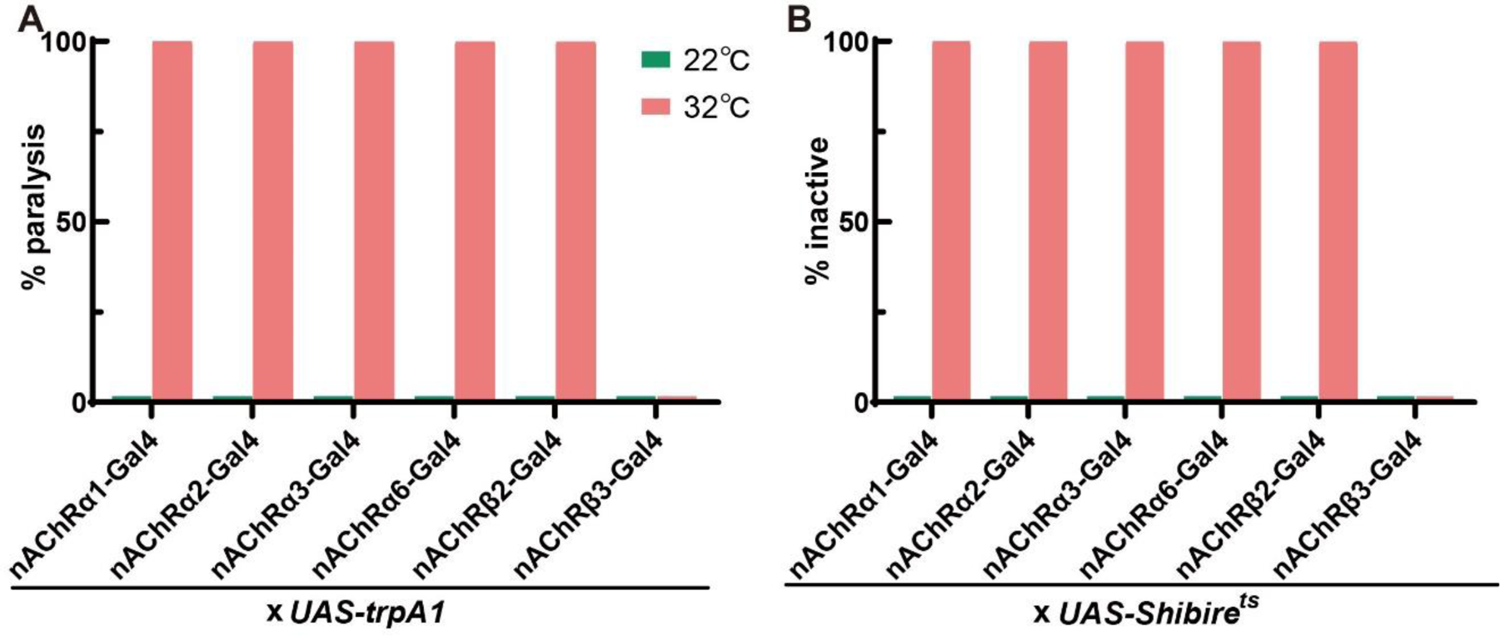
The effects of artificial neuronal activation and inhibition in various *nAChR*-expressing neurons. (A) Thermogenetic activation of five *nAChR*-expressing neurons using *UAS-trpA1* induced paralysis behavior. (B) Thermogenetic silencing of five *nAChR*-expressing neurons using *UAS-Shibire*^ts^ decreased activity. n = 30-50.

The poisoning symptoms associated with triflumezopyrim is distinct from other nicotinic modulation insecticides since it inhibits rather than activates insect nAChRs. There is no any neuro-excitatory symptoms after treatment of triflumezopyrim, on the contrary, it induces lethargic poisoning characterized by slow but coordinated leg movements and insects became less responsive to stimuli over time [19]. Thus, we chose to use *UAS-Shibire^ts^* to inhibit *nAChR-expressing* neurons [20]. As expected, *nAChRα1^2A-GAL4^*-*nAChRα2^2A-GAL4^ and nAChRβ2^2A-GAL4^* neurons produced a “sluggish” behavior rather than hyperactivity (Figure 3B). The flies exhibited almost no translational or rotational body movement (Video 1). Silencing of *nAChRα3^2A-GAL4^* and *nAChRα6^2A-GAL4^* neurons also produced similar behaviors, further confirming that the α3- and α6-containing nAChRs can not be blocked by triflumezopyrim, otherwise both mutants would show resistance in bioassays.

### Expression patterns of nAChRs in KO mutants

We confirmed that the KO coding regions were not detected or barely detectable with real-time PCR quantification (Figure S2). There was no big difference of expression levels of each subunits in these mutant flies, except that the β3 was relatively less transcribed than other genes. For the *α*1 heterozygous mutant, the mRNA levels of all subunits were almost same as the wild type control.

## Discussion

The Insecticide Resistance Action Committee (IRAC) classifies neonicotinoids, sulfoximines, butenolides and mesoionics into sub-groups 4A, 4C, 4D and 4E respectively, according to their chemical similarity relations. However, our results clearly showed that sulfoxaflor and flupyradifurone may specifically act on the same nAChR subtype which consists of α3 and β1 subunits (Figure 4A), albeit their big differences in chemical structures. More importantly, we found that the neonicotinoids act on distinct nAChR subtypes and such selectivity is not dependent on the aromatic heterocyclic (A), or the electron-withdrawing nitro or cyano moiety (X-Y) which is considered the key toxophore. Interestingly, the ring systems and the R_2_ substituents in the open-chain structures are the determining factors (Figure 4). For example, the α1, α2, β1 and β2 mutants showed similar levels of resistance to imidacloprid and thiacloprid (both have a five-membered ring), indicating that they mainly act on the same α1/α2/β1/β2 pentamer (Figure 4B). This is consistent with previous ex vivo recording results [21] and the two recent reconstituted studies in which both drugs act as partial agonists on the α1/α2/β1/β2 nAChR [7, 8]. Acetamiprid is structurally similar to thiacloprid with the cyanoimine phamacophore, but the acyclic configuration changes its molecular target in vivo. It may act on the α1/β1/β2 nAChR and again the electrophysiological studies had already indicated that acetamiprid is nearly a full agonist [21] and its potency on the recombinant louse α1/α2/β1/β2 nAChR is about 860 fold lower than that of thiacloprid [8]. Although thiamethoxam has a six-membered-ring, it is a pro-drug without intrinsic nAChR activity until metabolized to the active form clothianidine in plants and insects [22]. Therefore, thiamethoxam, clothianidine, dinotefuran and nitenpyram can be considered as the same type which have the N-methy substitution in the R_2_ position and mainly act on the α1/α3/β1 nAChR (Figure 4B). The neonicotinoids are traditionally divided into nitroimines (NNO_2_), nitromethylenes (CHNO2) or cyanoimines (NCN), but our findings proposed a new classification according to their major nAChR subtypes targets.

**Figure 4.**
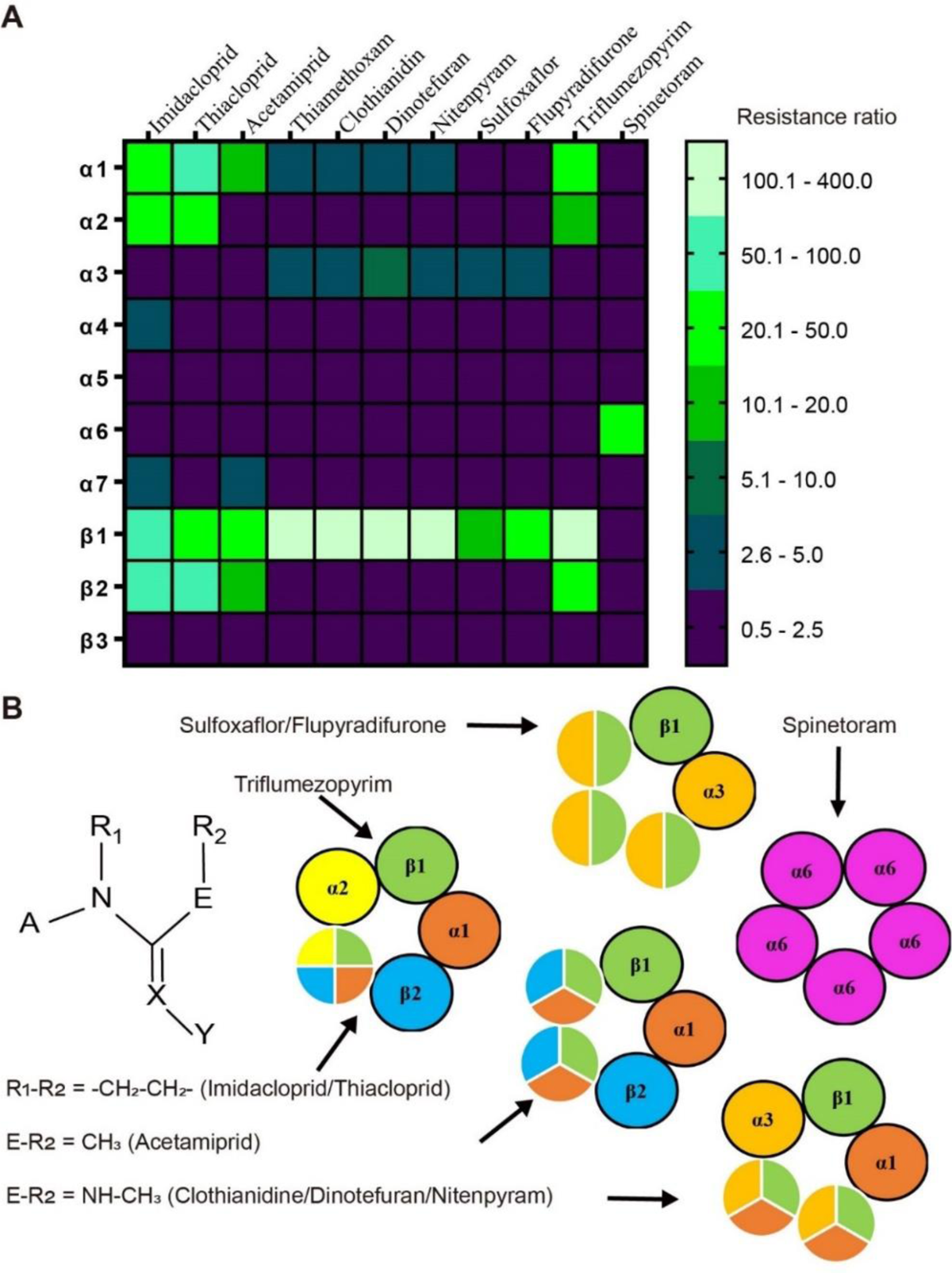
The resistance patterns of tested insecticides on different nAChR mutants (A) and the proposed target receptor subtypes for neonicotinoids and others (B). Various resistance ratios are grouped and represented as different colors in the heatmap. Thiamethoxam is considered as a prodrug of clothianidine and not listed in the structural formula.

Despite the widespread use of neonicotinoids for almost four decades, the first and only field-evolved target-site resistance mutation (R81T in nAChRβ1) was reported in 2011 and it has only been found in two species to date [11, 12]. Such unusual phenomenon can be partially explained by our findings that the seven neonicotinoids have at least three distinct molecular targets in vivo. To some extent, the continuous use of different neonicotinoids is a kind of spontaneous insecticides rotations, which has been proven to be effective in mitigating or delaying resistance. New nicotine mimic insecticides like sulfoxaflor and flupyradifurone mainly act on another nAChR subtype which is distinct from neonicotinoids (Figure 4), indicating their potential use in insecticides resistance management.

Electrophysiological studies with native tissues or recombinant receptors showed that low concentrations of neonicotinoids can block nAChR while higher concentrations cause receptor activation[7, 23]. Therefore, it is still unclear whether the insecticidal activity is the consequence of nAChR inhibition or activation in vivo. We found that transient artificial activation rather than inhibition of *nAChR-*expressing neuron is sufficient to induce neonicotinoids-like poisoning symptoms in flies (Figure 3). Thus, the overall effect of neonicotinoids is neuronal depolarizing by activation of nAChR which is more physiologically relevant.

Triflumezopyrim is the first member of a new class of mesoionic insecticides, which act via inhibition of the orthosteric binding site of the nAChR [19]. We found that the α1/α2/β1/β2 nAChR could be its major target like imidacloprid and thiacloprid, all these mutants showed high resistance to triflumezopyrim (Figure 4A). This is consistent with radioligand binding results in which triflumezopyrim potently displacing [^3^H]imidacloprid with a Ki value of 43 nM using the membrane preparations from the aphid [19]. Thermogenetic inhibition neurons expressing α1, α2 and β2 also mimic the lethargic intoxication symptoms (Figure 3B). Thus, in order to maintain the durability and effectiveness of this new powerful tool for control of hopper species in rice, it is very critical to avoid repeated use of triflumezopyrim with imidacloprid and thiacloprid.

The spinosyns including spinosad and spinoteram have been shown to act a population of nAChR that are not targeted by neonicotinoids, the binding site is also distinct to the orthosteric site [4]. The α6 subunit has been proposed as the main target of spinosyns since the resistance to spinosad in many insects is associated with loss-of-function mutations in the α6 gene [24], however, whether other subunits are involved is still unknown. We used spinoteram in bioassays and the results strongly indicated that spinosyns may specifically act on the α6 homomeric nAChR but not any other subtypes (Figure 4), which is consistent with a recent report using spinosad [25]. Thermogenetic activation of *α6-* expressing neurons also induced spinosyns-like poisoning symptoms in flies.

Our current knowledge about the subunit composition of insect nAChRs is very limited. Immunoprecipitation data with subunit-specific antibodies showed that the Drosophila *α*3 and β1 co-assemble within the same receptor complex [26]. Further studies from the same group indicated that *α*1/*α*2/β2 and β1/β2 may co-assemble into the same receptor complex respectively [27]. Similar studies using the brown planthopper suggested that there are two populations of nAChRs which contain the Drosophila equivalent subunits combinations *α*1/*α*2/β1 and *α*3/β1/β2, respectively [28]. These previous findings are partially confirmed by the present results, as the α3/β1, α1/α3/β1, α1/β1/β2 and α1/α2/β1/β2 could be the major receptor subtypes for the tested insecticides, indicating that the β1 subunit could be an indispensable component for all heteromecic pentamers (Figure 4). Besides, we noticed that for some insecticides, different subunits mutations contribute in an asymmetrical manner to resistance (Figure 4A). Therefore, there could be functional redundancy between some *α*-type subunits and we can not exclude the exitance of other potential receptor subtypes such as α1/β1 and α3/β1/β2. The diversity of insect nAChRs and their druggability make them remain an extremely important target for insecticides development.

Growing evidence indicates that sublethal doses of neonicotinoids like imidacloprid, thiamethoxam and clothianidin negatively affect wild and managed bees which are important pollinators in ecosystems and agriculture [29–31]. They reduce reproduction and colony development, perhaps by impairing foraging, homing and nursing behaviors of bees [32]. These severe sublethal effects have led to heavy restrictions on the use of above three neonicotinoids in Europe to protect pollinators [33]. The sulfoxaflor and flupyradifurone are potential alternatives for neonicotinoids, however, their risk to bees is controversial [34–36]. Therefore, it is critical to understand the mode of action of these insecticides inside bees. Since most *Drosophila* nAChR subunit genes (except *α*5 and β3) have one-to-one orthologs in the honeybee and bumblebee genomes [7], the expression and assembly of receptors could be conserved between flies and bees, suggesting that our results will enable further studies about the ecotoxicology and risk assessment for these nAChR modulators.

## Materials and Methods

### Insecticides

Imidacloprid (600g/LSC, Bayer CropScience, Germany), thiamethoxam (70%GZ, Syngenta, China), clothianidin (48%SC, HeNan Hansi crop protection, China), dinotefuran (20%SG, Mitsui Chemicals, Japan), nitenpyram (30%WG, ZinGrow, China), acetamiprid (20%SP, Noposion, China), thiacloprid (40%SC, Limin Chemical, China), sulfoxaflor (22%SC, Dow AgroSciences, USA), flupyradifurone (17%SC, Bayer CropScience, Germany), triflumezopyrim (10%SC, DuPont, USA), spinetoram (60g/LSC, Dow AgroSciences, USA) and triton X-100 (Sangon Biotech, China) were purchased commercially.

### Fly strains

Flies were maintained and reared on conventional cornmeal-agar-molasses medium at 25 ± 1 ℃, 60% ± 10% humidity with a photoperiod of 12 hours light: 12 hours night. For experiments using *UAS-trpA1* and *UAS-Shibire^ts^* transgenes, flies were reared at 21 ℃. The following stains were sourced from the Bloomington Stock Center (Indiana University): vas-cas (#51323), *UAS-trpA1* (#26263), *UAS-Shibire^ts^* (44222). All nAChR KO mutants and KI-Gal4 strains were gifts from Dr. Yi Rao (Deng et al., 2019) (Peking University). The *w^1118^* used for outcrossing was used as wide-type for insecticide bioassays.

We generated the nAChRβ1^R81T^ mutant by CRISPR/Cas9 genome editing. The gRNA sequence (3L:4433329∼4433352, ATCAAACGTTTGGTTAACTTTAG) was designed with flyCRISPR Target Finder (https://flycrispr.org/target-finder/) and cloned into the pDCC6 plasmid (addgene #59985). A 110 bp ssODN (single-strand oligodeoxynucleotide) was customer synthesized as the donor template to replace the targeted genomic region. This ssODN contained three nucleotides changes with two (CG to AC) conferring the R81T mutation and one synonymous mutation (G to A) to prevent the re-cleavage from Cas9 after incorporation. Both gRNA plasmid and ssODN were microinjected into the embryos of *vas-cas* flies (BL #51323). The crossing and selection scheme was shown in the Figure S1.

### Insecticide bioassays

3–5 day old and uniform size adult females were used in insecticide bioassays to assess the susceptibility of different fly strains. The testing method was modified from the IRAC susceptibility test method 026 (https://irac-online.org/methods/).

Briefly, the required serial dilutions of insecticide solution are prepared in 200g/L sucrose using formulated insecticides. Approximately 5ml of insecticide solution is required for each concentration. A piece of dental wick (1cm) is placed in a standard Drosophila vials and treated with 800 μL 20% aqueous sucrose with or without insecticide. The vials were kept upside down until all flies became active to avoid flies getting trapped in the dental wick. The bioassay was assessed after 48 h, dead flies as well as seriously affected flies displaying no coordinated movement, that were unable to walk up the vial, or unable to get to their feet were cumulatively scored as ‘affected’. The lethal concentrations LC_50_ were calculated by probit analysis using the Polo Plus software (LeOra Software, Berkeley, CA, USA). Non-linear log dose-response curves were generated in Graphpad Prism 8.21 (Graphpad Software Inc., La Jolla, CA, USA).

### Thermogenetic activation and silencing assays

Flies for TRPA1-mediated thermogenetic activation and Shibire-mediated silencing experiments were collected upon eclosion and reared in vials containing standard food medium at 21 ℃ for 5-8 days. For thermogenetic activation with the *UAS-trpA1* transgene, 10 flies were transferred to new empty vials by gently inspiration, and then the assays were performed at 23 ℃ and 32 ℃ for 10 minutes. The percentage of paralysis behavior, in which the animal lies on its back with little effective movement of the legs and wings, was measured. For silencing assays, *UAS-Shibire^ts^* transgene was used and flies were also transferred to fly vials at 23 ℃ and 32 ℃ for 10 minutes.

### Real-time quantitative PCR

The relative transcription levels of *nAChRs* in different KO mutants were examined using real-time quantitative PCR performed with an CFX96TM Real-Time PCR System (Bio-rad, Hercules, USA). Total RNA was isolated with Trizol reagent according to the manufacturer’s instructions. Residual genomic DNA was removed by RQ1 RNase-Free DNase (Promega). Total RNA was reverse transcribed to cDNA with the EasyScript First-Strand cDNA Synthesis SuperMix (Transgene, Beijing, China). qPCR with gene-specific primers was performed with the ChamQ Universal SYBR qPCR Master Mix (Vazyme, Nanjing, China) to investigate relative expression levels of different *nAChRs*. The *RpL32* (ribosomal protein L32) was used as an internal control. Relative expression of *nAChRs* were normalized to the reference (RpL32) using the 2^-ΔΔCT^ method.

### Fecundity and development assays

10 pairs of freshly emerged couples of wild type control and β1^R81T^ mutant were transferred into vials containing normal food for 72 hr. These files were then transferred into a new dish which is used for egg-laying assay. The numbers of egg laid in each dishes were recorded after 24 hr. To calculate the larvae to pupae rate, 60 second-instar larvae were collected and transferred into a new vial as one group. The numbers of pupae in each vial were recorded after 7 days in an incubator. Each genotypes were repeated for at least three times with duplicates.

### Climbing assay

About three-day-old male flies were collected with CO_2_ anesthesia into groups of 10, and then allowed to recover for 2 days. A climbing tube consisted of two vials with 90 mm height and 20 mm diameter. The flies were filmed for 30 s with a SONY HDR-CX900E camera. The climbing index (percentage of flies in the upper half of the vial) were determined at 5 s intervals, after the flies had been tapped down to the bottom of the vials.

## Acknowledgments

This work was supported by the National Natural Science Foundation of China (31802019, 32072496) and Zhejiang Provincial Fund for Distinguished Young Scholars (LR19C140002). We thank Yi Rao (Peking University) for fly stocks.

## Supplementary Information

**Figure S1.**
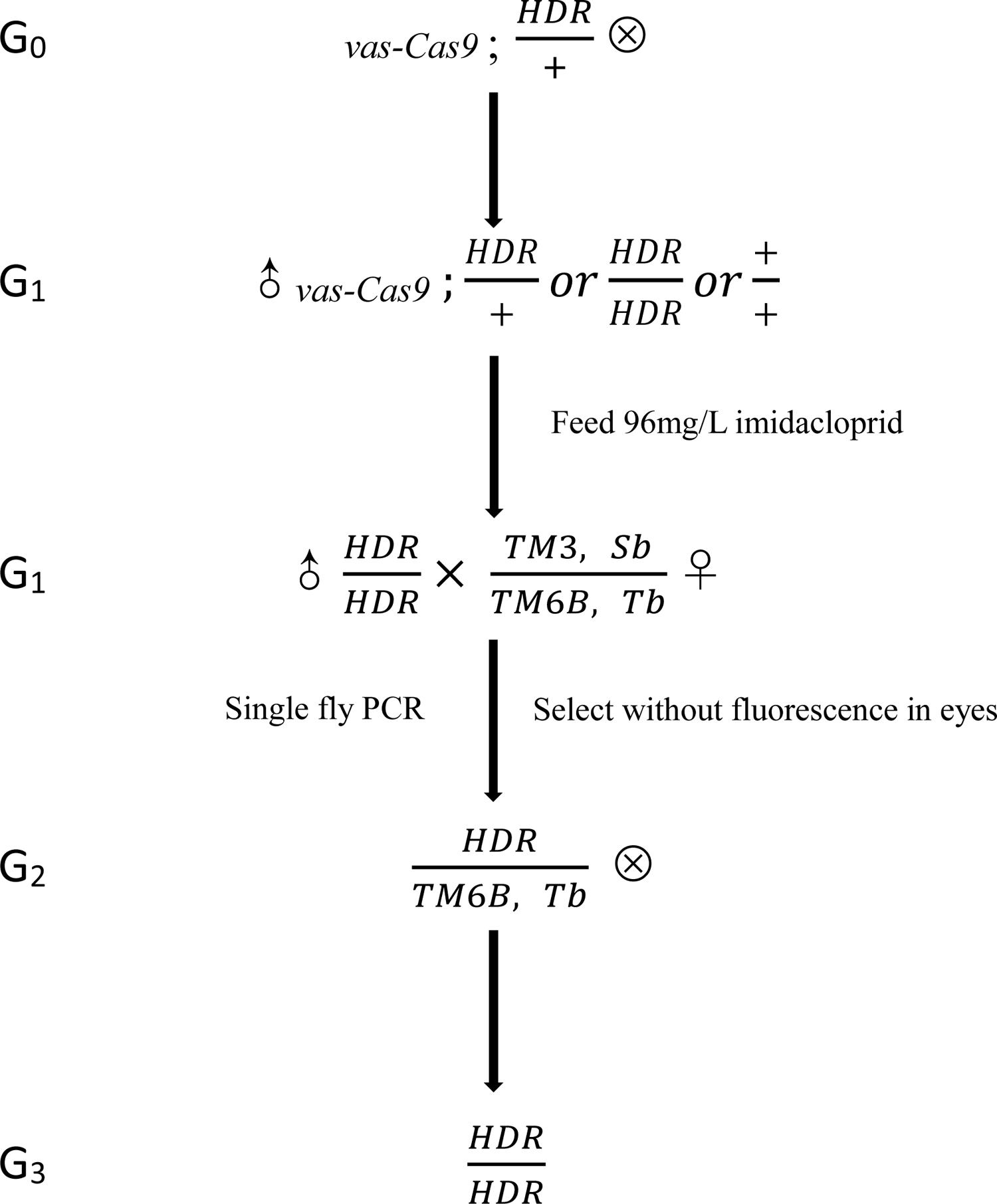
The crossing schemes to establish the *nAChRβ1^R81T^* knock-in line. The HDR event was isolated by imidacloprid selection and confirmed by PCR. The *vas-Cas9* (3XP3 RFP) was removed by the absence of red fluorescence in eyes.

**Figure S2.**
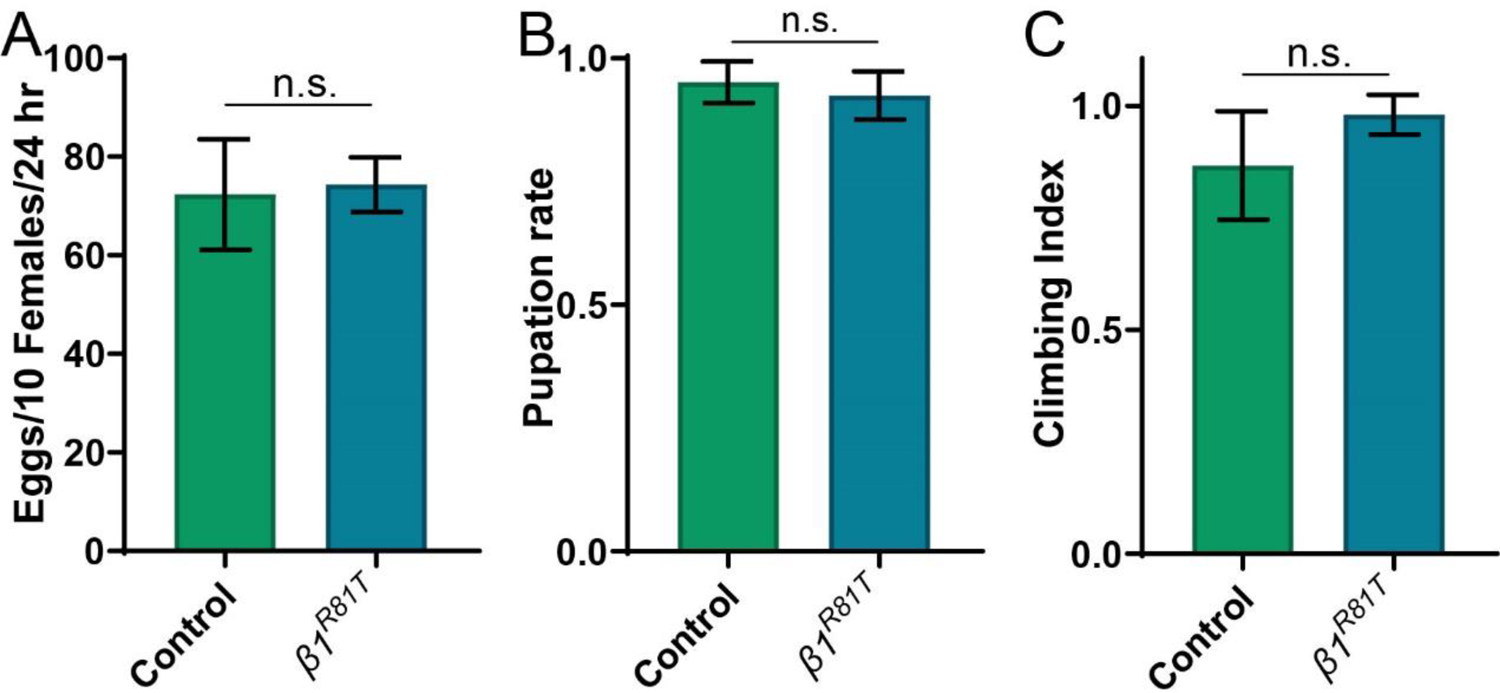
Effects of *nAChRβ1^R81T^* point mutation on number of eggs laid (A), pupation rate of larvae (B) and negative geotaxis behavior (C).

**Figure S3.**
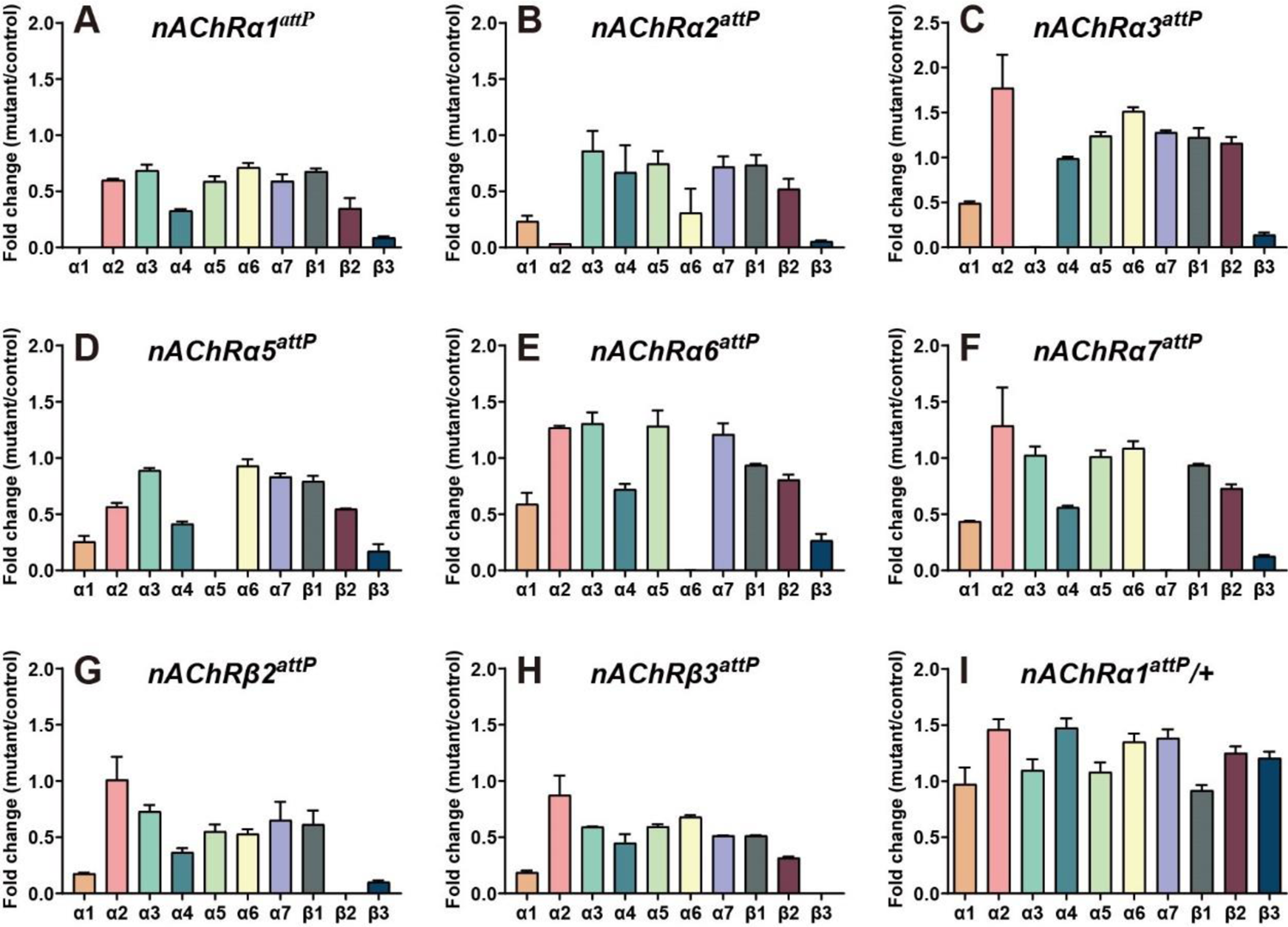
Expression patterns of the nAChR genes in different KO mutants.

**Video 1.** The effects of thermogenetic activation and inhibition in *nAChRα1-* expressing neurons. The following transgenes were used: *nAChRα1^2A-GAL4^* > *UAS-trpA1*; *nAChRα1^2A-GAL4^* > *UAS-Shibire^ts^*. Other *nAChR* KI-Gal4 strains like n*AChRα2^2A-GAL4^*, *nAChRα3^2A-GAL4^*, *nAChRα6^2A-GAL4^ and nAChRβ2^2A-GAL4^* also produced similar behaviors when stimulated under 32 ℃, these videos are not shown.

---

**Table S1.** Log dose probit mortality data and resistance ratios for imidacloprid

| Strain | LC<sub>50</sub><br>(mg/L) | 95% CL | LC<sub>95</sub><br>(mg/L) | 95% CL | Resistance ratio |  |
| --- | --- | --- | --- | --- | --- | --- |
|  |  |  |  |  | LC<sub>50</sub> | LC<sub>95</sub> |
| Control | 8.6 | 7.5-9.9 | 26.6 | 21.5-35.9 | 1.0 | 1.0 |
| $\alpha 1^{-/-}$ | 296.7 | 161.4-519.4 | 14555 | 5396.3-85114.0 | 34.5 | 547.2 |
| $\alpha 1^{+/-}$ | 150.4 | 74.7-272.7 | 935.5 | 456.0-5243.1 | 17.5 | 35.2 |
| $\alpha 2^{-/-}$ | 417.0 | 254.8-686.0 | 3113.5 | 1549.4-13500.0 | 48.5 | 117.0 |
| $\alpha 2^{+/-}$ | 52.5 | 38.7-69.6 | 114.0 | 82.2-259.1 | 6.1 | 4.3 |
| $\alpha 3^{-/-}$ | 10.7 | 9.7-11.7 | 16.7 | 14.7-21.1 | 1.2 | 0.6 |
| $\alpha 4^{T227M}$ | 22.9 | 19.2-27.2 | 58.0 | 45.2-86.9 | 2.7 | 2.2 |
| $\alpha 5^{-/-}$ | 9.6 | 7.5-12.1 | 27.5 | 19.9-50.2 | 1.1 | 1.0 |
| $\alpha 6^{-/-}$ | 7.8 | 6.2-10.0 | 28.2 | 19.9-49.7 | 0.9 | 1.1 |
| $\alpha 7^{-/-}$ | 22.8 | 20.4-25.5 | 43.6 | 37.1-55.8 | 2.7 | 1.6 |
| $\beta 1^{R81T}$ | 565.7 | 407.3-781.4 | 2885.7 | 1835.2-6036.1 | 65.8 | 108.5 |
| $\beta 1^{R81T/+}$ | 87.3 | 73.8-103.4 | 210.3 | 165.0-312.2 | 10.2 | 7.9 |
| $\beta 2^{-/-}$ | 725.4 | 513.6-1027.5 | 4702.2 | 2840.2-10564.0 | 84.3 | 176.8 |
| $\beta 2^{+/-}$ | 24.7 | 21.6-28.4 | 42.3 | 35.1-60.5 | 2.9 | 1.6 |
| $\beta 3^{-/-}$ | 13.1 | 11.6-14.7 | 27.5 | 23.1-35.6 | 1.5 | 1.0 |

**Table S2.** Log dose probit mortality data and resistance ratios for thiacloprid

| Strain | LC<sub>50</sub><br>(mg/L) | 95% CL | LC<sub>95</sub><br>(mg/L) | 95% CL | Resistance ratio |  |
| --- | --- | --- | --- | --- | --- | --- |
|  |  |  |  |  | LC<sub>50</sub> | LC<sub>95</sub> |
| Control | 30.4 | 28.3-32.6 | 49.3 | 44.6-56.7 | 1.0 | 1.0 |
| $\alpha I^{-/-}$ | 2674.9 | 2007.8-3553.3 | 8940.1 | 6082.5-17994.0 | 88.0 | 181.3 |
| $\alpha I^{+/-}$ | 89.2 | 67.1-116.2 | 182.2 | 134.3-394.2 | 2.9 | 3.7 |
| $\alpha 2^{-/-}$ | 1031.0 | 783.7-1384.3 | 14812.0 | 8693.3-31445.0 | 33.9 | 300.4 |
| $\alpha 2^{+/-}$ | 37.3 | 31.6-43.9 | 86.2 | 68.2-126.3 | 1.2 | 1.7 |
| $\alpha 3^{-/-}$ | 36.9 | 31.0-43.8 | 93.4 | 73.1-138.0 | 1.2 | 1.9 |
| $\alpha 4^{T227M}$ | 27.0 | 16.7-40.5 | 106.2 | 63.4-357.9 | 0.9 | 2.2 |
| $\alpha 5^{-/-}$ | 20.0 | 16.9-23.6 | 46.7 | 36.9-68.4 | 0.7 | 0.9 |
| $\alpha 6^{-/-}$ | 26.1 | 22.6-29.9 | 78.5 | 63.8-105.1 | 0.9 | 1.6 |
| $\alpha 7^{-/-}$ | 40.2 | 33.7-47.5 | 95.3 | 75.1-142.7 | 1.3 | 1.9 |
| $\beta I^{R81T}$ | 1342.7 | 802.5-2229.7 | 8533.6 | 4418.4-32285.0 | 44.2 | 173.1 |
| $\beta I^{R81T/+}$ | 40.3 | 26.3-62.2 | 106.9 | 67.4-458.1 | 1.3 | 2.2 |
| $\beta 2^{-/-}$ | 1677.8 | 922.8-3425.5 | 21015.0 | 8759.4-98473.0 | 54.3 | 401.8 |
| $\beta 2^{+/-}$ | 11.8 | 8.7-14.4 | 32.5 | 24.6-58.3 | 0.4 | 0.7 |
| $\beta 3^{-/-}$ | 39.4 | 33.5-46.4 | 88.0 | 70.0-128.4 | 1.3 | 1.8 |

**Table S3.** Log dose probit mortality data and resistance ratios for acetamiprid

| Strain | LC<sub>50</sub><br>(mg/L) | 95% CL | LC<sub>95</sub><br>(mg/L) | 95% CL | Resistance ratio |  |
| --- | --- | --- | --- | --- | --- | --- |
|  |  |  |  |  | LC<sub>50</sub> | LC<sub>95</sub> |
| Control | 9.7 | 7.4-13.1 | 22.0 | 15.5-50.7 | 1.0 | 1.0 |
| $\alpha I^{-/-}$ | 131.3 | 118.0-146.2 | 239.4 | 204.7-306.1 | 13.5 | 10.9 |
| $\alpha I^{+/-}$ | 43.6 | 31.0-59.2 | 90.2 | 64.7-240.8 | 4.5 | 4.1 |
| $\alpha 2^{-/-}$ | 15.2 | 11.4-20.5 | 37.9 | 26.2-89.1 | 1.6 | 1.7 |
| $\alpha 3^{-/-}$ | 16.9 | 14.7-19.5 | 43.1 | 34.5-61.8 | 1.7 | 2.0 |
| $\alpha 4^{T227M}$ | 9.4 | 8.2-10.9 | 17.3 | 14.2-24.8 | 1.0 | 0.8 |
| $\alpha 5^{-/-}$ | 10.1 | 8.8-11.5 | 23.9 | 19.7-32.0 | 1.0 | 1.1 |
| $\alpha 6^{-/-}$ | 22.6 | 18.5-28.0 | 63.7 | 46.6-109.0 | 2.3 | 2.9 |
| $\alpha 7^{-/-}$ | 27.4 | 18.8-41.2 | 55.3 | 38.0-204.6 | 2.8 | 2.5 |
| $\beta I^{R81T}$ | 231.5 | 214.7-249.5 | 397.7 | 355.5-465.5 | 23.9 | 18.1 |
| $\beta I^{R81T/+}$ | 31.2 | 27.4-35.4 | 50.5 | 43.0-67.6 | 3.2 | 2.3 |
| $\beta 2^{-/-}$ | 126.5 | 109.6-146.4 | 238.3 | 195.2-340.1 | 13.0 | 10.8 |
| $\beta 2^{+/-}$ | 24.9 | 22.1-28.3 | 38.0 | 32.4-51.8 | 2.6 | 1.7 |
| $\beta 3^{-/-}$ | 17.7 | 13.5-23.3 | 55.6 | 38.2-111.7 | 1.8 | 2.5 |

**Table S4.** Log dose probit mortality data and resistance ratios for thiamethoxam

| Strain | LC<sub>50</sub>(mg/L) | 95% CL | LC<sub>95</sub>(mg/L) | 95% CL | Resistance ratio |  |
| --- | --- | --- | --- | --- | --- | --- |
|  |  |  |  |  | LC<sub>50</sub> | LC<sub>95</sub> |
| Control | 8.2 | 7.6-8.9 | 15.3 | 13.4-18.4 | 1.0 | 1.0 |
| $\alpha 1^{-/-}$ | 30.5 | 22.7-41.7 | 65.8 | 46.6-158.5 | 3.7 | 4.3 |
| $\alpha 2^{-/-}$ | 7.9 | 6.8-9.2 | 14.9 | 12.2-21.2 | 1.0 | 1.0 |
| $\alpha 3^{-/-}$ | 32.9 | 29.5-36.4 | 71.2 | 61.6-87.1 | 4.0 | 4.7 |
| $\alpha 4^{T227M}$ | 4.1 | 2.6-5.9 | 9.9 | 6.6-34.8 | 0.5 | 0.6 |
| $\alpha 5^{-/-}$ | 7.3 | 6.6-8.0 | 11.3 | 10.0-13.6 | 0.9 | 0.7 |
| $\alpha 6^{-/-}$ | 10.8 | 10.0-11.6 | 16.7 | 14.8-20.4 | 1.3 | 1.1 |
| $\alpha 7^{-/-}$ | 8.2 | 7.2-9.3 | 16.5 | 13.7-22.1 | 1.0 | 1.1 |
| $\beta 1^{R81T}$ | 1935.5 | 1756.0-2142.6 | 4783.9 | 4047.9-5980.8 | 236.0 | 312.7 |
| $\beta 1^{R81T/+}$ | 18.3 | 12.9-25.0 | 38.4 | 27.4-97.2 | 2.2 | 2.5 |
| $\beta 2^{-/-}$ | 12.8 | 11.0-14.9 | 25.8 | 20.9-37.0 | 1.6 | 1.7 |
| $\beta 3^{-/-}$ | 11.8 | 8.9-20.0 | 22.6 | 15.4-121.0 | 1.4 | 1.5 |

**Table S5.** Log dose probit mortality data and resistance ratios for clothianidin

| Strain | LC<sub>50</sub><br>(mg/L) | 95% CL | LC<sub>95</sub><br>(mg/L) | 95% CL | Resistance ratio |  |
| --- | --- | --- | --- | --- | --- | --- |
|  |  |  |  |  | LC<sub>50</sub> | LC<sub>95</sub> |
| Control | 3.5 | 3.1-3.9 | 8.4 | 7.0-10.9 | 1.0 | 1.0 |
| $\alpha 1^{-/-}$ | 10.3 | 9.2-11.4 | 20.5 | 17.5-26.1 | 2.9 | 2.4 |
| $\alpha 2^{-/-}$ | 4.7 | 4.3-5.2 | 8.5 | 7.4-10.5 | 1.3 | 1.0 |
| $\alpha 3^{-/-}$ | 11.5 | 8.1-16.7 | 25.9 | 17.5-78.1 | 3.3 | 3.1 |
| $\alpha 4^{T227M}$ | 2.5 | 2.1-2.9 | 6.3 | 4.9-9.5 | 0.7 | 0.8 |
| $\alpha 5^{-/-}$ | 2.8 | 2.5-3.2 | 4.4 | 3.8-5.7 | 0.8 | 0.5 |
| $\alpha 6^{-/-}$ | 5.1 | 4.6-5.7 | 9.4 | 8.1-11.9 | 1.5 | 1.1 |
| $\alpha 7^{-/-}$ | 6.1 | 5.3-6.9 | 17.3 | 14.1-23.2 | 1.7 | 2.1 |
| $\beta 1^{R81T}$ | 969.4 | 730.6-1278.8 | 4940.0 | 3300.4-9182.5 | 277.0 | 588.1 |
| $\beta 1^{R81T}/+$ | 13.4 | 11.8-15.2 | 21.2 | 18.1-28.7 | 3.8 | 2.5 |
| $\beta 2^{-/-}$ | 3.4 | 3.0-4.0 | 6.2 | 5.2-8.6 | 1.0 | 0.7 |
| $\beta 3^{-/-}$ | 7.6 | 7.0-8.4 | 13.5 | 11.7-16.9 | 2.2 | 1.6 |

**Table S6.** Log dose probit mortality data and resistance ratios for dinotefuran

| Strain | LC<sub>50</sub><br>(mg/L) | 95% CL | LC<sub>95</sub><br>(mg/L) | 95% CL | Resistance ratio |  |
| --- | --- | --- | --- | --- | --- | --- |
|  |  |  |  |  | LC<sub>50</sub> | LC<sub>95</sub> |
| Control | 5.5 | 5.2-5.9 | 11.4 | 10.2-13.0 | 1.0 | 1.0 |
| $\alpha 1^{-/-}$ | 15.0 | 13.6-17.0 | 25.9 | 21.7-34.8 | 2.7 | 2.3 |
| $\alpha 2^{-/-}$ | 7.2 | 6.5-7.9 | 12.3 | 10.7-15.4 | 1.3 | 1.1 |
| $\alpha 3^{-/-}$ | 30.4 | 27.6-33.2 | 55.0 | 48.4-66.0 | 5.5 | 4.8 |
| $\alpha 3^{+/-}$ | 16.2 | 14.1-18.7 | 29.2 | 24.2-41.3 | 2.9 | 2.6 |
| $\alpha 4^{T227M}$ | 3.7 | 3.2-4.3 | 7.0 | 5.7-10.2 | 0.7 | 0.6 |
| $\alpha 5^{-/-}$ | 6.5 | 6.0-7.2 | 9.7 | 8.6-11.8 | 1.2 | 0.9 |
| $\alpha 6^{-/-}$ | 8.7 | 7.8-9.8 | 16.7 | 14.1-21.7 | 1.6 | 1.5 |
| $\alpha 7^{-/-}$ | 7.3 | 6.7-8.0 | 10.5 | 9.5-12.2 | 1.3 | 0.9 |
| $\beta 1^{R81T}$ | 2190.9 | 1790.6-2702.1 | 4803.3 | 3659.6-7985.8 | 398.3 | 421.3 |
| $\beta 1^{R81T/+}$ | 9.3 | 8.1-10.6 | 15.6 | 13.0-22.2 | 1.7 | 1.4 |
| $\beta 2^{-/-}$ | 10.3 | 9.0-11.8 | 17.6 | 14.7-24.7 | 1.9 | 1.5 |
| $\beta 3^{-/-}$ | 11.4 | 8.0-20.6 | 23.7 | 15.4-201.0 | 2.1 | 2.1 |

**Table S7.** Log dose probit mortality data and resistance ratios for nitenpyram

| Strain | LC<sub>50</sub><br>(mg/L) | 95% CL | LC<sub>95</sub><br>(mg/L) | 95% CL | Resistance ratio |  |
| --- | --- | --- | --- | --- | --- | --- |
|  |  |  |  |  | LC<sub>50</sub> | LC<sub>95</sub> |
| Control | 19.8 | 17.9-22.2 | 36.6 | 30.8-48.9 | 1.0 | 1.0 |
| $\alpha I^{-/-}$ | 54.4 | 43.5-69.5 | 115.2 | 85.4-220.8 | 2.7 | 3.1 |
| $\alpha 2^{-/-}$ | 15.1 | 13.4-17.0 | 33.1 | 27.7-43.0 | 0.8 | 0.9 |
| $\alpha 3^{-/-}$ | 53.7 | 44.8-64.3 | 147.3 | 113.2-223.5 | 2.7 | 4.0 |
| $\alpha 4^{T227M}$ | 13.4 | 11.2-15.7 | 28.9 | 23.0-44.3 | 0.7 | 0.8 |
| $\alpha 5^{-/-}$ | 16.0 | 14.5-17.5 | 24.7 | 22.0-29.9 | 0.8 | 0.7 |
| $\alpha 6^{-/-}$ | 19.4 | 16.1-24.8 | 35.4 | 27.7-61.1 | 1.0 | 1.0 |
| $\alpha 7^{-/-}$ | 13.6 | 12.2-15.2 | 25.9 | 22.0-33.2 | 0.7 | 0.7 |
| $\beta I^{R8IT}$ | 3629.0 | 2906.8-4770.6 | 11394.0 | 7852.2-21070.0 | 183.3 | 311.3 |
| $\beta I^{R8IT/+}$ | 30.3 | 26.6-34.7 | 51.4 | 43.2-70.4 | 1.5 | 1.4 |
| $\beta 2^{-/-}$ | 29.6 | 25.8-34.0 | 52.2 | 43.4-72.6 | 1.5 | 1.4 |
| $\beta 3^{-/-}$ | 17.1 | 15.5-19.0 | 29.6 | 25.5-37.5 | 0.9 | 0.8 |

**Table S8.**
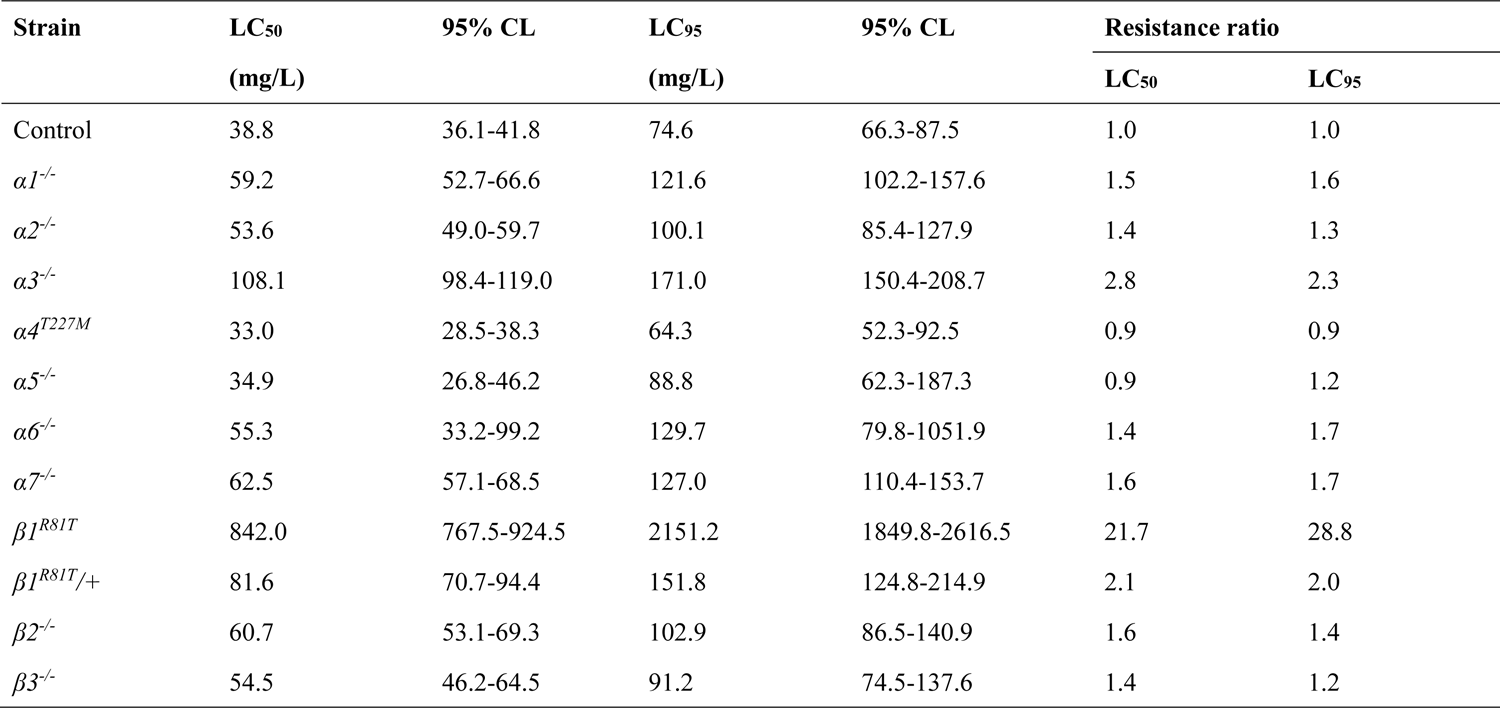
Log dose probit mortality data and resistance ratios for flupyradifurone

**Table S9.** Log dose probit mortality data and resistance ratios for sulfoxaflor

| Strain | LC<sub>50</sub><br>(mg/L) | 95% CL | LC<sub>95</sub><br>(mg/L) | 95% CL | Resistance ratio |  |
| --- | --- | --- | --- | --- | --- | --- |
|  |  |  |  |  | LC<sub>50</sub> | LC<sub>95</sub> |
| Control | 59.4 | 56.1-62.8 | 90.6 | 83.9-99.9 | 1.0 | 1.0 |
| $\alpha 1^{-/-}$ | 54.8 | 50.0-60.3 | 85.3 | 75.3-103.2 | 0.9 | 0.9 |
| $\alpha 2^{-/-}$ | 82.0 | 73.9-91.0 | 142.7 | 122.8-181.6 | 1.4 | 1.6 |
| $\alpha 3^{-/-}$ | 207.8 | 185.6-236.3 | 306.0 | 263.2-402.5 | 3.5 | 3.4 |
| $\alpha 4^{T227M}$ | 73.0 | 50.8-108.0 | 166.2 | 110.7-594.5 | 1.2 | 1.8 |
| $\alpha 5^{-/-}$ | 55.7 | 50.7-61.3 | 88.5 | 77.9-107.6 | 0.9 | 1.0 |
| $\alpha 6^{-/-}$ | 63.0 | 57.3-69.0 | 96.6 | 86.0-115.7 | 1.1 | 1.1 |
| $\alpha 7^{-/-}$ | 56.6 | 51.5-62.1 | 88.2 | 78.0-106.2 | 1.0 | 1.0 |
| $\beta 1^{R81T}$ | 948.5 | 726.2-1245.3 | 2280.2 | 1628.6-4643.7 | 16.0 | 25.2 |
| $\beta 1^{R81T/+}$ | 116.3 | 102.9-131.0 | 175.1 | 152.0-222.1 | 2.0 | 1.9 |
| $\beta 2^{-/-}$ | 50.3 | 46.4-54.8 | 76.2 | 67.6-91.7 | 0.8 | 0.8 |
| $\beta 3^{-/-}$ | 64.2 | 59.7-68.1 | 89.7 | 83.9-98.4 | 1.1 | 1.0 |

**Table S10.** Log dose probit mortality data and resistance ratios for triflumezopyrim

| Strain | LC<sub>50</sub><br>(mg/L) | 95% CL | LC<sub>95</sub><br>(mg/L) | 95% CL | Resistance ratio |  |
| --- | --- | --- | --- | --- | --- | --- |
|  |  |  |  |  | LC<sub>50</sub> | LC<sub>95</sub> |
| Control | 28.2 | 24.7-31.9 | 237.0 | 184.0-328.0 | 1.0 | 1.0 |
| $\alpha I^{-/-}$ | 922.1 | 756.1-1066.0 | 12856.0 | 9343.9-19626.0 | 32.7 | 54.2 |
| $\alpha 2^{-/-}$ | 484.4 | 266.6-985.3 | 13397.0 | 3076.9-71026.0 | 17.2 | 56.5 |
| $\alpha 3^{-/-}$ | 36.0 | 24.7-47.1 | 448.5 | 278.0-1005.9 | 1.3 | 1.9 |
| $\alpha 4^{T227M}$ | 18.2 | 13.8-22.8 | 73.4 | 51.4-141.1 | 0.6 | 0.3 |
| $\alpha 5^{-/-}$ | 18.0 | 1.0-15.2 | 43.3 | 34.2-63.5 | 0.6 | 0.2 |
| $\alpha 6^{-/-}$ | 28.7 | 24.3-34.0 | 142.5 | 105.2-219.3 | 1.0 | 0.6 |
| $\alpha 7^{-/-}$ | 30.0 | 24.5-37.0 | 238.1 | 157.0-440.0 | 1.1 | 1.0 |
| $\beta I^{R81T}$ | 4349.2 | 3096.0-6661.3 | 33601 | 17279.0-117310.0 | 154.0 | 141.8 |
| $\beta 2^{-/-}$ | 668.9 | 431.0-913.5 | 5797.4 | 3728.1-12256.0 | 23.7 | 24.5 |
| $\beta 3^{-/-}$ | 28.2 | 24.7-31.9 | 237.0 | 184.0-327.9 | 1.5 | 0.7 |

**Table S11.** Log dose probit mortality data and resistance ratios for spinetora

| Strain | LC<sub>50</sub><br>(mg/L) | 95% CL | LC<sub>95</sub><br>(mg/L) | 95% CL | Resistance ratio |  |
| --- | --- | --- | --- | --- | --- | --- |
|  |  |  |  |  | LC<sub>50</sub> | LC<sub>95</sub> |
| Control | 1.3 | 1.2-1.4 | 2.5 | 2.2-3.1 | 1.0 | 1.0 |
| $\alpha 1^{-/-}$ | 1.8 | 1.2-2.6 | 4.0 | 2.7-13.5 | 1.4 | 1.6 |
| $\alpha 2^{-/-}$ | 0.8 | 0.7-0.9 | 1.8 | 1.5-2.3 | 0.6 | 0.7 |
| $\alpha 3^{-/-}$ | 1.2 | 1.0-1.3 | 2.6 | 2.2-3.5 | 0.9 | 1.0 |
| $\alpha 4^{T227M}$ | 1.8 | 1.3-2.5 | 4.4 | 3.0-11.1 | 1.4 | 1.8 |
| $\alpha 5^{-/-}$ | 1.4 | 1.2-1.6 | 4.0 | 3.2-5.6 | 1.1 | 1.6 |
| $\alpha 6^{-/-}$ | 55.6 | 45.8-68.1 | 134.3 | 101.8-218.1 | 42.8 | 53.7 |
| $\alpha 6^{+/-}$ | 1.6 | 1.4-1.8 | 3.4 | 2.9-4.5 | 1.2 | 1.4 |
| $\alpha 7^{-/-}$ | 1.8 | 1.6-2.2 | 4.8 | 3.8-6.7 | 1.4 | 1.9 |
| $\beta 1^{R81T}$ | 0.8 | 0.7-0.9 | 2.1 | 1.7-2.8 | 0.6 | 0.8 |
| $\beta 2^{-/-}$ | 0.8 | 0.7-0.9 | 1.3 | 1.1-1.7 | 0.6 | 0.5 |
| $\beta 3^{-/-}$ | 1.4 | 1.2-1.6 | 2.9 | 2.4-3.8 | 1.1 | 1.2 |
| $\alpha 5^{-/-}; \alpha 7^{-/-}$ | 2.2 | 1.8-2.5 | 4.5 | 3.6-6.6 | 1.7 | 1.8 |

**Table S12.**
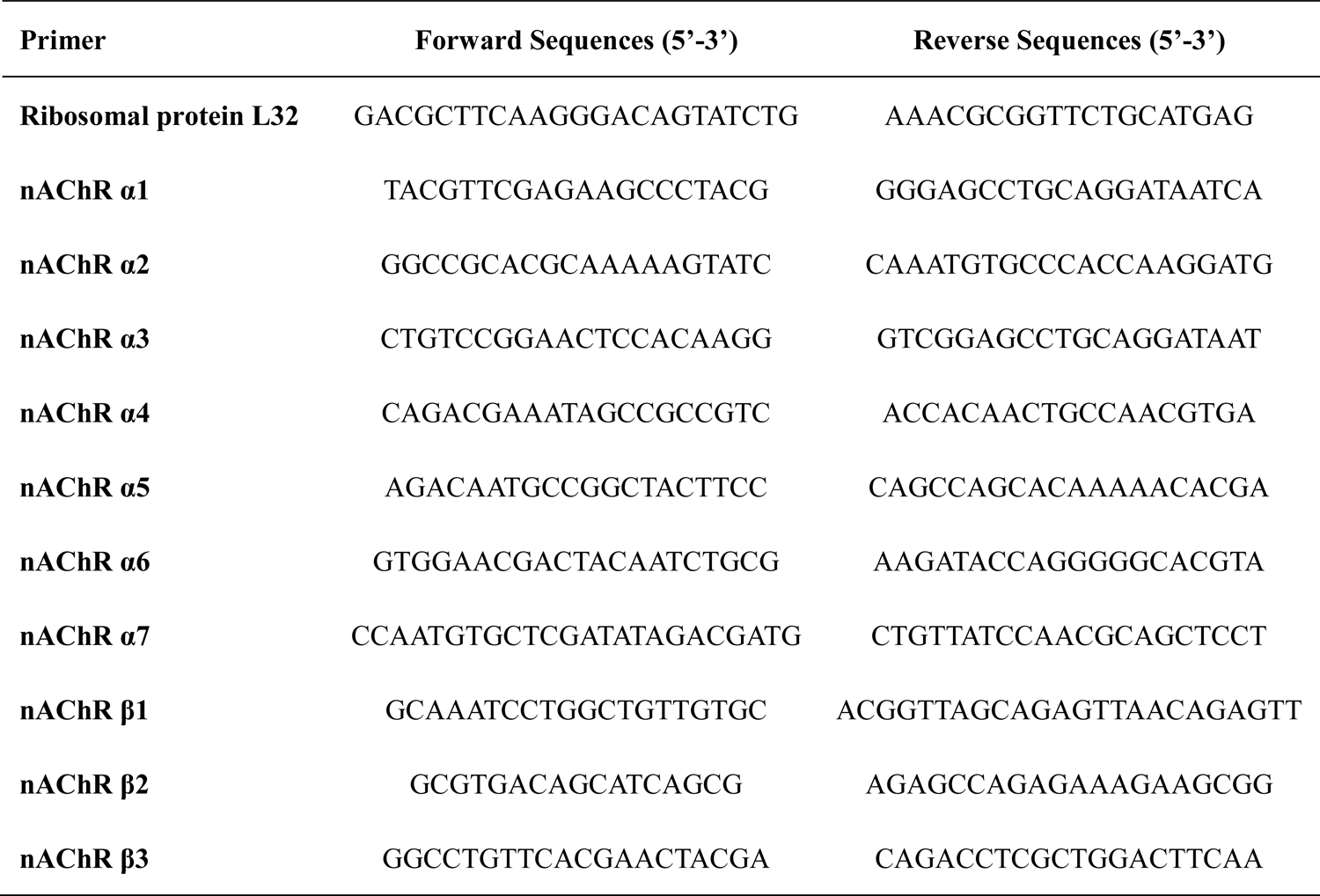
Primers used in qPCR analysis

